# TBI-166 inhibits DlaT and LpdC, exhibiting synergistic effects with Bedaquiline and Pyrazinamide against *Mycobacterium tuberculosis*

**DOI:** 10.64898/2026.05.20.726459

**Authors:** Chennan Liu, Lei Zhang, Zimo Wang, Xiaodong Li, Bin Wang, Yu Lu

## Abstract

Tuberculosis (TB), particularly drug-resistant tuberculosis (DR-TB), remains a critical global challenge and underscores the urgent need for novel drugs and innovative combination regimens with distinct mechanisms of action. Here, we characterize an all-oral three-drug regimen comprising TBI-166, bedaquiline(BDQ), and pyrazinamide(PZA), which displays strong synergistic antimicrobial activity in vitro against both replicating and non-replicating *Mycobacterium tuberculosis* (MTB) and has previously shown superior bactericidal and sterilizing efficacy to standard HRZ and BPaL regimens in murine TB models(1). Time-kill studies demonstrate that the triple regimen outperforms dual-drug combinations, accelerating bacterial clearance across multiple physiological states.

Mechanistic investigations revealed that the TBI-166–BDQ–PZA combination induces a comprehensive collapse of energy and redox homeostasis, marked by profound ATP depletion, robust accumulation of reactive oxygen species (ROS), and marked disruption of the intracellular NAD(H) pool. TBI-166, a novel riminophenazine analogue of clofazimine (CFZ) currently in phase II clinical trials, emerged as a key contributor to this metabolic stress. Metabolomic profiling and ¹³C-based flux analysis show that TBI-166 slows glycolysis and the tricarboxylic acid (TCA) cycle while enhancing flux through the pentose phosphate and nicotinate pathways, thereby lowering the NADH/NAD⁺ ratio and diminishing MTB metabolic flexibility under environmental stress. In parallel, TBI-166 downregulates the dormancy regulator DosR and its regulon, further compromising adaptation to non-replicating states.

Multi-omics analyses, together with biochemical and biophysical assays, identify the pyruvate dehydrogenase complex (PDHc) components DlaT and LpdC as direct molecular targets of TBI-166, with drug binding leading to potent inhibition of their enzymatic activities. Collectively, these findings define the mechanism of action of TBI-166 and provide a molecular rationale for its inclusion in potent, all-oral, short-course regimens. More broadly, they highlight the therapeutic potential of metabolically targeted combinations that destabilize energy metabolism, redox balance, and metabolic adaptability to improve DR-TB treatment outcomes.

## 1. Introduction

Despite the advances in medical interventions, drug-resistant tuberculosis (DR-TB) remains a global health challenge, with the World Health Organization (WHO) reporting approximately 150,000 deaths attributed to DR-TB in 2023 (2). The limitations of traditional antibiotics exacerbate this challenge, as they predominantly target actively replicating bacteria, leaving unaffected non-replicating populations. This severely undermines treatment effectiveness and promotes the development of resistant strains (3). In addition, the remarkable metabolic adaptability of *Mycobacterium tuberculosis* (MTB) makes treatments more complex. The bacterium can rapidly switch between different physiological states, enabling survival under diverse and hostile environmental conditions, making complete eradication extremely difficult (4, 5). This metabolic plasticity allows MTB to evade conventional antibiotic treatments, contributing to treatment failure and high relapse rates (6, 7). Moreover, only 68% of DR-TB patients successfully complete treatment (2), underscoring the urgent need for novel drugs and innovative therapeutic approaches, particularly those that can target MTB across multiple metabolic states (8). Building on our previous work demonstrating that an all-oral three-drug regimen comprising TBI-166, Bedaquiline (BDQ), and Pyrazinamide (PZA), achieves superior bactericidal and sterilizing activity to standard HRZ and BPaL regimens in murine TB models(1), we sought to characterize the mechanistic basis of this enhanced efficacy. In these models, the TBI-166+BDQ+PZA combination demonstrated excellent sterilizing activity, enhanced early bactericidal effects, and significantly reduced relapse rates, highlighting its promising potential for treating DR-TB(1).

Despite this encouraging in vivo performance, the precise mechanisms underlying the enhanced therapeutic efficacy of this three-drug combination remain incompletely understood. BDQ disrupts ATP synthesis by binding to the c and ε subunits of ATP synthase, uncoupling the electron transport chain (ETC) from energy production and leading to the accumulation of toxic metabolic intermediates (9–13). PZA, activated to pyrazinoic acid (POA) by pncA, disrupts membrane potential and causes a depletion of energy production (14, 15). TBI-166 is a structural optimization of clofazimine (CFZ), maintains comparable antimicrobial activity while significantly minimizing skin discoloration side effects, with its primary mechanism associated with disrupting the redox balance of MTB (16–20). Building upon these mechanistic insights, we hypothesized that the synergistic effects of the TBI-166+BDQ+PZA combination effectively address MTB’s metabolic plasticity through comprehensive metabolic inhibition across various bacterial physiological states. However, the specific contribution of TBI-166 and its molecular targets remain unclear. Building upon these unresolved questions, this study aimed to systematically investigate the synergistic effects of TBI-166, BDQ, and PZA across different MTB physiological states. Concurrently, we aimed to conduct an in-depth exploration of TBI-166’s molecular mechanism, focusing on identifying its specific molecular targets and unique contributions to metabolic disruption in MTB. Our primary objectives were to: (1) evaluate the growth inhibition potential of the three-drug combination against replicating and non-replicating MTB; (2) characterize the initial inhibitory mechanisms; and (3) elucidate the specific role of TBI-166 in this multi-targeted therapeutic approach. By comprehensively examining these aspects, we sought to provide mechanistic insights into the potential of this novel drug combination for more effective tuberculosis treatment.

## 2. Materials and methods

### 2.1 Bacterial Strains and Culture Conditions

*M. tuberculosis* H37Rv (ATCC 27294) was grown in 7H9 broth (Difco, BD271310) supplemented with 0.05% Tween 80 and 10% GOADC enrichment medium PLUS (Gene-Optimal, GOMY0078). AR-MTB cultures were incubated at 37°C with 5% CO_2_. NR-MTB was generated using two established models: the Wayne hypoxic model and nutrient starvation model. CFU was analyzed as previously described (1).

### 2.2 Chemicals

TBI-166(MIC=0.06μg/mL) was provided by the Institute of Materia Medica & Peking Union Medical College (Beijing, China); BDQ(MIC=0.04μg/mL), PZA was purchased from Sigma-Aldrich. All other chemicals were of analytical grade and obtained from Sigma-Aldrich unless otherwise specified.

### 2.3 Biochemical Assays

Intracellular ATP levels were measured using an ATP Assay Kit (Beyotime, S0026) according to the manufacturer’s instructions. Briefly, bacterial cells were harvested by centrifugation (8000g, 4℃, 10 min), and the pellet was resuspended in 1 mL of substrate solution to achieve a final concentration of approximately 10^8^ CFU/mL. ATP content was determined as previously described (21).

Intracellular ROS and NAD(H) levels were measured using commercial assay kits. ROS levels were determined using a ROS Detection Kit (AAT Bioquest, 22903). Bacterial samples were incubated with the probe for 1 h, and fluorescence intensity was measured at excitation/emission wavelengths of 650/675 nm. NAD(H) content was assessed using a NAD(H) Assay Kit (AAT Bioquest, 15263). After 1 h probe incubation, fluorescence was monitored at excitation/emission wavelengths of 540/590 nm. All fluorescence measurements were performed using an Infinite 200 multimode microplate reader (Tecan, Männedorf, Switzerland).

### 2.4 RNA Sequencing and Analysis

Total RNA was extracted using a RNeasy Mini Kit (QIAGEN, 74524) according to the manufacturer’s instructions. RNA quality and quantity were assessed using a NanoDrop spectrophotometer and an Agilent BioAnalyzer 2100 system (Agilen Technologies.). RNA sequencing was performed by Bohao Biotechnology Co., Ltd. (Shanghai, China) using the Illumina platform. Briefly, following rRNA depletion, mRNA was fragmented and reverse transcribed to construct the cDNA library according to the Illumina mRNA Sequencing Sample Preparation Guide. Single-end sequencing was performed on an Illumina Genome Analyzer IIx using two separate lanes under the control of Illumina data collection software.

### 2.5 ^13^C Metabolic Flux Analysis

Metabolomic profiling was performed using 13C-labeled glucose as a metabolic tracer. D-Glucose-13C6 (MedChemExpress, HY-B0389A) was used for metabolic flux analysis. Log-phase H37Rv cells were washed three times with 1× PBS and resuspended in 10 mL 7H9 medium supplemented with either 0.5% 13C-labeled or unlabeled glucose. After 20 h incubation, bacterial cells (10^7 CFU) were quenched with 400 μL pre-chilled extraction solution (methanol:acetonitrile, 1:1, v/v). Quality control samples were prepared by pooling equal volumes of supernatant from unlabeled samples. Metabolite analysis was performed using a Vanquish UHPLC system coupled to an Orbitrap Exploris 120 mass spectrometer (Thermo Fisher Scientific). Polar metabolites were separated on a Waters ACQUITY UPLC BEH Amide column (2.1 × 100 mm, 1.7 μm) using 25 mM ammonium acetate and 25 mM ammonium hydroxide in water (mobile phase A) and acetonitrile (mobile phase B). Non-polar metabolites were separated on a Phenomenex Kinetex C18 column (2.1 × 100 mm, 2.6 μm) using 0.01% acetic acid in water (mobile phase A) and isopropanol:acetonitrile (1:1, v/v) (mobile phase B), Mass spectrometry was performed using Xcalibur software (version 4.4).

### 2.6 Metabolomics Analysis

Metabolites were extracted from drug-treated bacterial cells (10^8 CFU) after washing three times with pre-chilled 1× PBS. Cells were resuspended in 1 mL pre-chilled methanol and stored at −80°C for 4 h. After spin-drying, samples were reconstituted in 300 μL water and subjected to three freeze-thaw cycles between liquid nitrogen and dry ice, followed by vortexing (30 s) and sonication (15 min) in an ice-water bath. A 250 μL aliquot of the supernatant was mixed with 750 μL pre-chilled methanol (−40°C), with the remaining supernatant reserved for protein quantification. After vortexing and incubation at −40°C for 1 h, samples were centrifuged (13,800 × g, 4°C, 15 min). The supernatant (900 μL) was collected, dried, and reconstituted in 180 μL ultrapure water before filtration for HPIC-MS/MS analysis.

Standard solutions were prepared by diluting individual compounds to 10 mmol/L stock solutions, followed by serial dilutions to create calibration curves. Chromatographic separation was performed on a Thermo Scientific Dionex ICS-6000 HPIC System equipped with Dionex IonPac AS11-HC (2 × 250 mm) and AG11-HC (2 × 50 mm) columns.Mass spectrometry was performed using an AB SCIEX 6500 QTRAP+ triple quadrupole mass spectrometer with ESI interface. Calibration curves were constructed using 1/x weighted least squares regression, with acceptance criteria of 80-120% accuracy. Detection limits (LOD and LOQ) were determined using signal-to-noise ratios of 3 and 10, respectively, following US FDA guidelines. Method validation included precision (relative standard deviation) and accuracy (analytical recovery) assessments using quality control samples.

### 2.7 Proteomics Analysis

Protein extraction was performed following drug treatment. Briefly, samples were precipitated with 4 volumes of pre-chilled 10% TCA-acetone at −20°C for 3 h. After centrifugation, the pellet was washed three times with 4 volumes of pre-chilled pure acetone (−20°C, 30 min), followed by centrifugation at 20,000 × g for 30 min at 4°C. The pellet was then dissolved in lysis buffer (8 M urea, 30 mM HEPES, 1 mM PMSF, 2 mM EDTA, and 10 mM DTT) and sonicated for 5 min (180 W power, 2 s pulse-on/3 s pulse-off). The lysate was clarified by centrifugation at 20,000 × g for 30 min. Protein reduction was performed by adding DTT to a final concentration of 10 mM and incubating at 56°C for 1 h, followed by alkylation with 55 mM iodoacetamide (IAM) in darkness for 1 h. Protein concentration was determined using the Bradford assay. Subsequent mass spectrometry analysis and proteomic data processing were performed by BGI (Shanghai, China). Data acquisition and analysis were conducted using Skyline (version 3.7.0.11317), Proteome Discoverer (version 1.4, Thermo Fisher Scientific), and SEQUEST HT.

### 2.8 Molecular Docking

Molecular docking was performed using MOE software to investigate protein-ligand interactions. The crystal structures of DlaT (Rv2215; PDB ID: 9Y7V), DlaT at the trimer-trimer interface (PDB ID: 9Y6T), and LpdC (PDB ID: 7KMY) were used as receptor models for docking analysis. Specifically, docking was conducted in the CoA-binding pocket of DlaT using 9Y7V, at the trimer-trimer interface of DlaT using 9Y6T, and in the ligand-binding region of LpdC using 7KMY. The Triangle Matcher algorithm was employed as the placement method with the London δG scoring function for initial pose evaluation. The top 1,000 poses were refined using the Induced Fit method and scored with the GBVI/WSA scoring function. The final 100 poses were clustered based on protein-ligand interaction fingerprints. The best-scoring pose was selected for subsequent analysis, considering various energy terms including conformational energy (E_Conf), placement energy (E_Place), and comprehensive scoring values (E_Score1 and E_refine).

### 2.9 Genetic Validation and Statistical Analysis

Overexpression strains of DlaT (Rv2215) and LpdC (Rv0462) were constructed using the pMV261 plasmid vector. The coding sequences were cloned into pMV261 via EcoRI and HindIII restriction sites, and recombinant plasmids were electroporated into H37Rv competent cells. Transformants were selected on Middlebrook 7H10 agar plates containing kanamycin (50 μg/mL) and validated by colony PCR and qRT-PCR. Knockdown strains in M. smegmatis were generated using the pGrna-MS0903 (MS4283) plasmid expressing target-specific gRNA in combination with the pTet-dCAS9 plasmid expressing dCas9 under tetracycline control, with knockdown efficiency confirmed by qRT-PCR(22). For M. tuberculosis knockdown strains, the PLJR965 CRISPRi plasmid was employed, expressing dCas9 and target-specific gRNA under anhydrotetracycline (ATc)-inducible control. Strains were selected on Middlebrook 7H10 agar plates containing hygromycin (50 μg/mL), with gene silencing efficiency validated by qRT-PCR after ATc (50 ng/mL) induction. Drug susceptibility testing was performed using the microplate Alamar Blue assay. Minimum inhibitory concentrations (MICs) of TBI-166 were determined for wild-type, overexpression, and knockdown strains.

## 3. Result

### 3.1 Synergistic Bactericidal Activity of the TBI-166+Bedaquiline+Pyrazinamide Combination via ATP Depletion and ROS Overproduction in Replicating *Mycobacterium tuberculosis*

We previously reported that the drug combination of TBI-166 + BDQ+ PZA exhibited synergistic bactericidal activity in H37Rv-infected C3HeB/FeJ and BALB/c mice, which may be related to their collective action on the energy synthesis pathway of MTB (1). To elucidate the mechanisms underlying the enhanced therapeutic efficacy of the three-drug combination and test our hypothesis of coordinated metabolic disruption, we conducted comprehensive bactericidal kinetic studies examining TBI-166, BDQ, and PZA both individually and in various combinatorial contexts.

Given that the in vitro activity assay of PZA needs to be conducted under acidic conditions (23), the bactericidal kinetics of each drug were first examined at different pH levels. At pH 7.0, TBI-166 exhibited growth inhibition at 1/4×MIC (0.015 µg/mL), 5×MIC (0.3 µg/mL), and 80×MIC (5 µg/mL); BDQ demonstrated bacteriostatic effects at 1/2×MIC (0.015 µg/mL) and bactericidal effects above its MIC; and PZA showed no bacteriostatic activity at any concentration tested. At pH 6.0, high concentrations of TBI-166 and PZA exhibited bactericidal activity. However, at pH 5.0, Mycobacterium tuberculosis viability was severely compromised (Figure S1). Therefore, subsequent synergy studies using the triple drug combination were conducted at pH 7.0 to allow for more robust observation of synergistic effects.

Under pH 7.0 conditions, the triple combination of TBI-166, BDQ, and PZA exhibited potent synergistic bactericidal activity against replicating H37Rv.At sub-inhibitory concentrations, the triple combination of TBI-166 (1/2×MIC), BDQ (1/2×MIC), and PZA (50 µg/mL) achieved a ∼4 log₁₀ CFU reduction after 7 days of treatment, an effect that substantially exceeded that of any single agent and surpassed the TBI-166+BDQ and TBI-166+PZA dual combinations by roughly 3 and 2 log₁₀ units, respectively (Figure 1A).Time-kill kinetic analysis over 14 days at bactericidal concentrations provided further resolution of this synergy: the triple combination (TBI-166 32×MIC + BDQ 8×MIC + PZA 100 µg/mL) sterilized the culture by day 7, whereas the TBI-166+BDQ dual combination required 10–14 days to reach the limit of detection, and the TBI-166+PZA dual combination failed to achieve sterilization within the 14-day observation window, with residual CFU counts remaining approximately 3 log₁₀ above the detection limit.(Figure 1B). Crucially, these kinetic data reveal a clear functional hierarchy—TBI-166+BDQ+PZA > TBI-166+BDQ > TBI-166+PZA—and establish a mechanistic division of labor: BDQ serves as the primary bactericidal engine, PZA accelerates the rate of BDQ-mediated killing, and TBI-166 amplifies the bactericidal depth and speed achieved by both partners.

**Figure 1.**
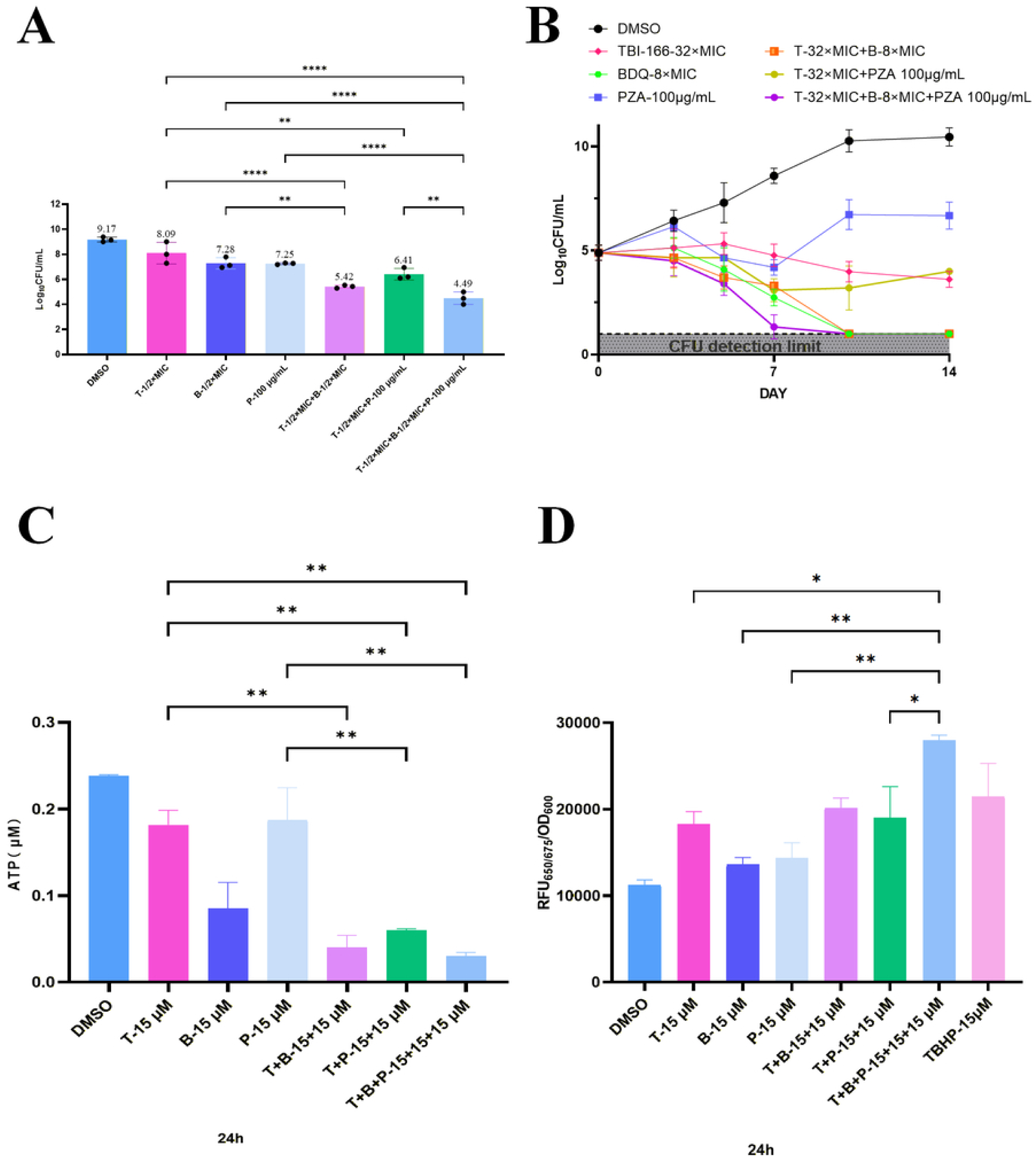
TBI-166 combined with BDQ and PZA exhibits potent synergistic bactericidal activity and triggers comprehensive collapse of energy and redox homeostasis in *Mycobacterium tuberculosis.* (A)CFU counts of *replicating Mycobacterium tuberculosis* H37Rv after 7 days of treatment with indicated sub-inhibitory concentrations ofTBI-166. BDQ, PZA, and their combinations. (B)Time-kill curves of replicating H37Rv treated with the highly concentrationsofTBI-166.BDQ,and PZA over 14 days. The grey shaded area represents the limit of detection. (C-D) The triple combination exacerbates the disruption of bioenergetic and redox homeostasis in MTB. evidenced by profound intracellular ATP depletion (C) and robust reactive owgen species(ROS) accumulation (D). TBHP was used as a positive control for ROS induction in (D).T: TBI-166: B: BDQ: P: PZA: TBHP: tert-Butyl hydroperoxicle. *:p<0.05; **: p<0.0L ***p<0.00L ****: p<0.0001.

We next sought to investigate the metabolic basis underlying this hierarchy. BDQ exerts its anti-MTB activity primarily through selective inhibition of the F₀F₁-ATP synthase, thereby disrupting ATP synthesis and impairing bacterial energy production (9–13), and PZA has also been reported to disturb the energy production of MTB (24); and TBI-166, as a structural analog of CFZ, is known to interfere with redox homeostasis through NADH-dependent redox cycling (19). Based on these distinct yet complementary mechanisms, we hypothesized that the synergistic superiority of the triple combination—particularly its ability to accelerate BDQ-mediated killing—derives from the coordinated disruption of both energy metabolism and redox balance.

To test this hypothesis, we assessed intracellular ATP levels in H37Rv following 24 h of drug exposure. Consistent with BDQ’s established role as the primary ATP synthase inhibitor, ATP levels were most profoundly reduced in regimens containing BDQ. TBI-166 alone (32×MIC) produced a significant but partial reduction in ATP compared with the untreated control, whereas BDQ alone (8×MIC) caused a more pronounced depletion. The TBI-166+BDQ dual combination reduced ATP to a level comparable to that of BDQ alone, and the triple combination (TBI-166 32×MIC + BDQ 8×MIC + PZA 100 µg/mL) drove ATP to nearly undetectable levels (Figure 1C). Notably, however, the ATP levels in the triple combination and the TBI-166+BDQ dual combination were statistically indistinguishable, despite the markedly superior bactericidal activity of the triple regimen observed in time-kill assays (Figure 1B). This dissociation between the degree of ATP depletion and the extent of bacterial killing suggested that ATP starvation alone could not fully account for the enhanced activity of the triple combination over the dual-drug regimens.

We therefore examined reactive oxygen species (ROS) as an additional contributor to cell death. TBI-166 alone at 32×MIC induced a significant increase in intracellular ROS levels, confirming that ROS generation is an intrinsic property of this clofazimine analog (Figure 1D). The TBI-166+BDQ dual combination further elevated ROS levels above those of either single agent. However, it was the triple combination that triggered a massive and sustained accumulation of intracellular ROS, reaching levels significantly higher than those induced by any single agent or dual-drug combination and, notably, even exceeding the effect of the positive control tert-butyl hydroperoxide (TBHP) (Figure 1D). Critically, the ROS level in the triple combination was significantly higher than that in the TBI-166+BDQ dual combination, providing the clearest biochemical parameter that distinguishes these two regimens and precisely mirrors their divergent killing kinetics (Figure 1B).

Taken together, these data demonstrate that the synergistic bactericidal superiority of the triple combination resides in a coordinated dual assault on MTB physiology: severe ATP depletion driven primarily by BDQ-mediated F₀F₁-ATP synthase inhibition, and overwhelming oxidative stress driven by TBI-166 and amplified by the energetic collapse induced by BDQ and PZA. While ATP depletion alone is comparable between the triple and TBI-166+BDQ dual combinations, the addition of PZA in the triple regimen markedly amplifies ROS production to levels that correlate with the accelerated and deepened killing observed in the time-kill assays. Thus, it is the convergence of profound energy deprivation and extreme redox imbalance—rather than either mechanism in isolation—that provides the mechanistic basis for the superior synergistic bactericidal efficacy of the TBI-166+BDQ+PZA regimen.

### 3.2 TBI-166 Disrupts NAD(H) Homeostasis and Reprograms Embden–Meyerhof–Parnas and Tricarboxylic Acid Cycle Activity in *Mycobacterium tuberculosis*

Given that TBI-166 served as the primary driver of ROS production in the triple combination, we next sought to dissect its intracellular mechanism of action. A central clue lies in the NAD(H) redox couple, which governs both electron flux through the respiratory chain and the generation of ROS via redox cycling (25, 26). We therefore investigated the impact of TBI-166 on NAD(H) homeostasis in H37Rv. Treatment with TBI-166 for 24 h resulted in a significant reduction in total NAD(H) pool (Figure 2A) and a marked decrease in the NADH/NAD⁺ ratio (Figure 2B) compared to the untreated control. Notably, NADH levels were more profoundly depleted than NAD⁺ levels, suggesting the existence of compensatory mechanisms that partially restore NAD⁺ under stress conditions. The precise molecular target through which TBI-166 elicits these effects remains to be fully elucidated. However, based on the well-established mechanism of its parent scaffold—CFZ, a phenazine compound known to undergo spontaneous redox cycling by competing with menaquinone for binding to NDH-2 dehydrogenase, thereby driving NADH oxidation and diverting electrons to generate ROS(25, 27)—it is plausible that TBI-166, as a structurally optimized phenazine analog, may perturb NAD(H) homeostasis through a similar mode of action. Nevertheless, given the structural differences introduced during optimization, the exact binding site and molecular interactions of TBI-166 within the mycobacterial target require further investigation.

**Figure 2.**
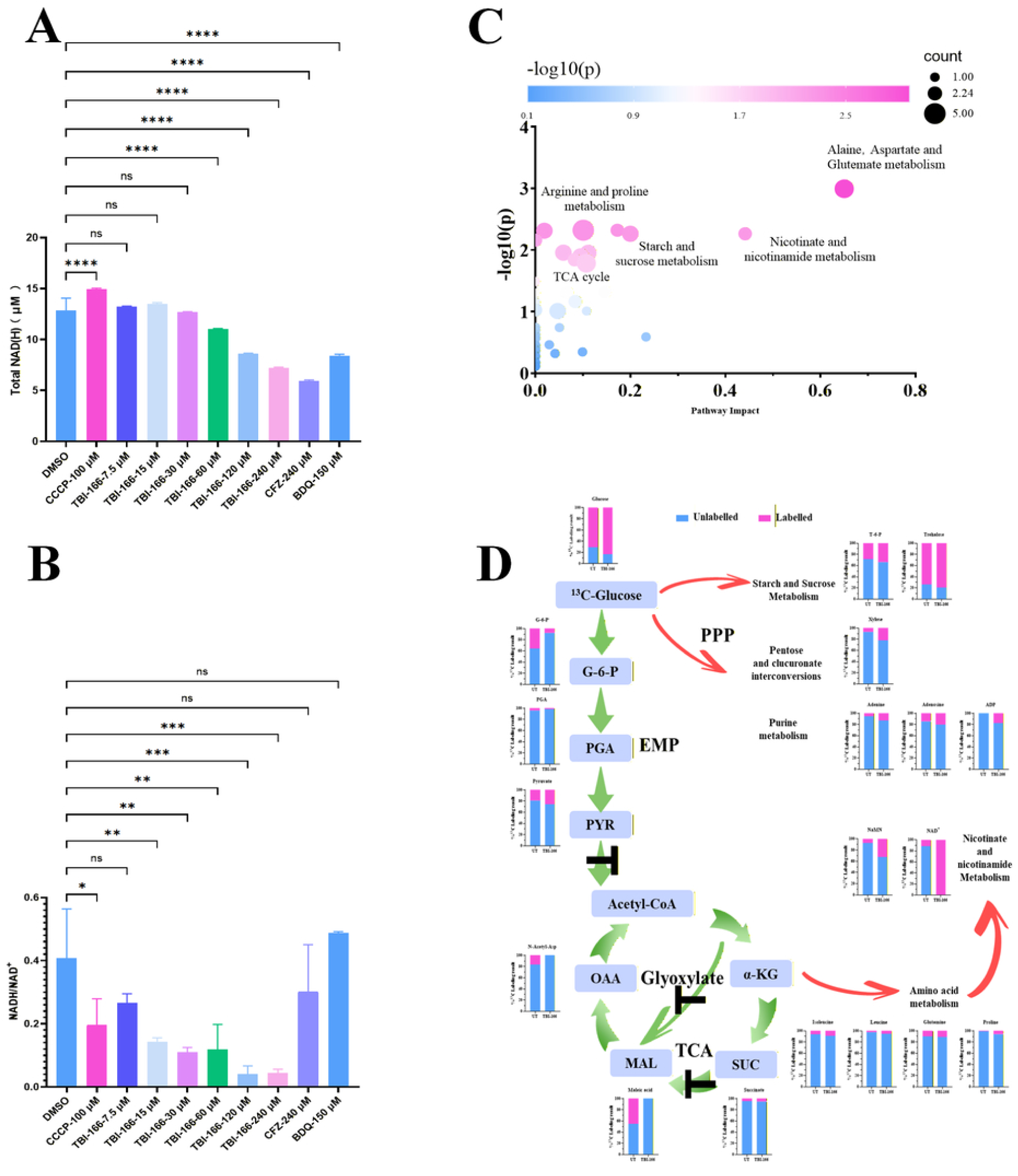
TBI-166 depletes the NAD(H) pool and reprograms central carbon metabolism in *Mycobacterium tuberculosis*. (A) Eftects of clilforent treatments on the total NAD(H) pool in MTB. Treatment with TBI-166 (7.5 pM to 240 µ1M) leads to a close--clepenclent depletion of total NAD(l-f) levels. (B) Eftects on the intracellular NADH/NAD+ ratio. The protonophore CCCP. CFZ. and BDQ are included as controls. (C) ^13^C isotopologue pathway enrichment analysis (Pathway Impact) of clifterentially abundant metabolites. The color gradient of the bubbles from blue to pink represents the decreasing p-value. with darker pink indicating higher statistical significance of the enrichment. The size of each bubble corresponds to the pathwav impact score. (D) Carbon isotopologue distribution and carbon flux in central carbon metabolic pathways. Bar charts illustrate the proportion of unlabeled (blue) and ^13^C-labeled (pink) carbons for specific metabolites in the untreated (UT) and TBI-166 treated 24h groups. The black T-bars:the inhibited metabolic steps: Reel arrows: increased metabolic flux: Green arrows: decreased metabolic flux. EMP, Embden-Meyerhof:Parnas pathway: PPP, pentose phosphate patlnYav: TCA. tricarboxylic acid cycle: G-6-P. glucose-6-phosphate: PGA, 3-phosphoglycerate: PYR, pyruvate: Acetyl-CoA. acetyl-coenzyme A: OAA. oxaloacetate: *o.-KG,a-*ketoglutarate:SUC, succinate: MAL malate: NaMN. nicotinate D-ribonucleotide: NAD+. nicotinamide adenine dinucleotide: N-Acetyl-Asp, N-acetyl-L-aspartate: T-6-P. trehalose-6-phosphate.

And the perturbation of NAD(H) homeostasis raised a further question: does TBI-166 also affect the metabolic pathways that produce and regenerate these redox cofactors? To address this, we performed pathway enrichment analysis and ¹³C-glucose tracing-based metabolomics to map the metabolic reprogramming induced by TBI-166 (5×MIC) in H37Rv. Untreated replicating MTB metabolizes ^13^C glucose via both glycolysis and gluconeogenesis, with glycolytic intermediates, such as pyruvate(PYR), feeding into the TCA cycle (28).Pathway enrichment analysis revealed that TBI-166 treatment significantly perturbed multiple metabolic pathways, with the most pronounced alterations occurring in the pentose phosphate pathway (PPP), the Embden Meyerhof Parnas (EMP) pathway, the tricarboxylic acid (TCA) cycle, and amino acid biosynthesis pathways (Figure 2C).

We then analyzed the ¹³C labeling patterns of central carbon metabolism intermediates to infer pathway-level flux alterations (Figure 2D). In this ¹³C-glucose tracing experiment, a higher ¹³C labeling fraction in each metabolite reflects a greater relative contribution of glucose-derived carbon to that metabolite pool, which may result from either increased flux from glucose or accumulation when downstream steps are blocked. Following TBI-166 treatment, the proportion of ¹³C-labeled glucose increased, consistent with active glucose uptake and limited dilution by unlabeled carbon sources. In contrast, ¹³C enrichment in the first EMP intermediates, glucose-6-phosphate (G-6-P) and 3-phosphoglycerate (PGA), was markedly reduced, whereas labeling of the pentose phosphate pathway (PPP) intermediate xylose and purine metabolites was increased. Together with the increased labeling of trehalose-6-phosphate (T-6-P) and trehalose in the starch and sucrose metabolism pathway(28, 29), these data indicate that the glycolytic (EMP) flux originating from glucose is attenuated, while PPP and trehalose biosynthesis become preferential routes for G-6-P utilization under TBI-166 exposure(Figure 2D).

Notably, despite the reduced labeling of upstream EMP intermediates, ¹³C incorporation into PYR, the end product of glycolysis, was increased. This apparent paradox does not reflect a more active EMP pathway; rather, it is best explained by a downstream block at the conversion of PYR to acetyl-CoA, which prevents PYR from entering the tricarboxylic acid (TCA) cycle and causes labeled PYR to accumulate potentially, reflecting ANA node regulation of EMP pathway intermediates and TCA cycle flux (29, 30). Consistent with an impaired TCA entry and progression, ¹³C labeling in malate (MAL) and oxaloacetate (OAA) dropped to undetectable levels, while succinate (SUC) labeling remained essentially unchanged. The simultaneous loss of MAL and OAA labeling, together with the absence of additional ¹³C incorporation beyond the succinate node, indicates that both the oxidative TCA cycle and the glyoxylate shunt are effectively blocked downstre(13)am of PYR(31, 32). In line with this, ¹³C labeling of multiple amino acids was increased, and NaMN as well as NAD exhibited enhanced ¹³C incorporation, suggesting that, once the TCA cycle is stalled, carbon from α-ketoglutarate is increasingly diverted into amino acid biosynthesis and ultimately into de novo NAD⁺ synthesis as a compensatory response to TBI-166–induced perturbation of NAD(H) homeostasis(Figure 2D).

### 3.3 TBI-166 Cooperates with Bedaquiline and Pyrazinamide to Disrupt Central Carbon Metabolism by Enhancing EMP/TCA Inhibition and Pentose Phosphate Pathway Flux in *Mycobacterium tuberculosis*

The above isotopologue-labelling data qualitatively demonstrate that TBI-166 reprograms MTB glucose metabolism, but ¹³C enrichment reflects relative carbon source usage rather than absolute metabolite abundance(13). We therefore quantitatively profiled central carbon metabolism (CCM) intermediates in MTB after TBI-166(5× MIC) treatment (Figure 3A). Consistent with the ¹³C tracing results, TBI-166 markedly increased the absolute levels of multiple pentose phosphate pathway (PPP) intermediates, including erythrose-4-phosphate (E4P), ribose-5-phosphate (R5P) and sedoheptulose-7-phosphate (S7P), as well as metabolites that sit at the EMP/PPP branch point, G6P, fructose-6-phosphate (F6P) and glyceraldehyde-3-phosphate (GAP). In parallel, several downstream glycolytic and TCA-linked metabolites, such as phosphoenolpyruvate (PEP), PYR, α-KG, SUC, MAL and fumarate (FUM), were also elevated, whereas the committed EMP intermediate fructose-1,6-bisphosphate (FBP), the triose phosphate glycerone-P (dihydroxyacetone phosphate) and citrate (CIT) were significantly decreased. Together, these data indicate that TBI-166 reduces net glycolytic (EMP) flux through the FBP and glycerone-P nodes, while promoting carbon accumulation at the G6P/F6P/GAP branch and within the PPP. The depletion of CIT, the first TCA intermediate generated from EMP-derived acetyl-CoA, further supports a restriction of carbon entry from glycolysis into the oxidative TCA cycle (Figure 2D).

**Figure 3.**
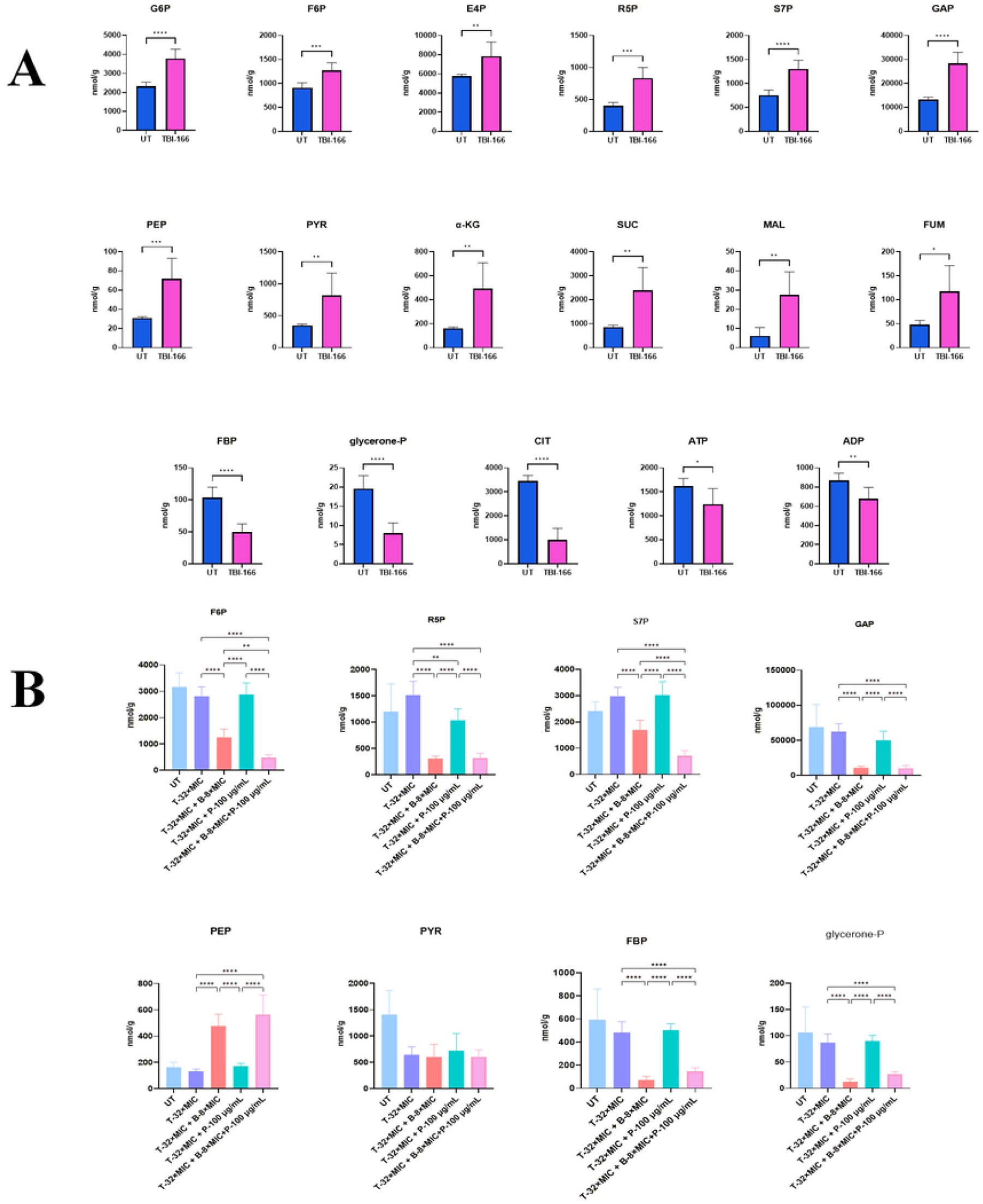
TBI-166 potentiates BDQ’s disruptive metabolism effects in *Mycobacterium tuberculosis.* leading to enhanced EMP/TCA cycle blockage and PPPintennediate accumulation, further amplified by PZA. (A)Quantification of central carbon metabolites in MTB after untreated(UT) or TBI-166 (5 XMIC) for 24 hours. Data are from six biologically independent replicates (n = 6). Statistical significance was determined by one-way ANOVA follo\Yecl by Dunnetfs multiple-comparison test. (B) Quantification of central carbon metabolites in MTB after untreated(UT,)TBI-166 (32 X MIC) or combination for 24 hours. Data are from nine biologicallv independent replicates (n = 9). Thecombinations included TBI-166 (T-32 XMIC) and BDQ (B-8X MIC) and/or PZA (P-100 1g/111L). EMP. Embden-Meyerhof-Parnas pathway: PPP. pentose phosphate pathway: TCA. tricarboxylic acid cvcle: G6P. glucose-6-phosphate: F6P. fructose-6-phosphate: E4P, ervthrose-4-phosphate: RSP. ribose-5-phosphate: S7P. sedoheptulose-7-phosphate 14: GAP, glyceraldehyde-3-phosphate 14: PEP. Phosphoenolpyruvate: PYR. pyruvate: u-KG. u-ketoglutarate: SUC. succinate: MAL. malate: FUM. fumarate: FBP. fructose-1.6-bisphosphate: glycerone-P. dihydroxyacetone phosphate: CIT. citrate: ATP. adenosine triphosphate: ADP. adenosine di phosphate. *:p<0.05; **: p<0.0 1; ***p<0.00 1: ****: p<0.0001.

To determine how this metabolic state interacts with BDQ and PZA, we next quantified the same CCM intermediates under single- and multi-drug conditions (Figure 3B). BDQ alone, consistent with previous work showing that ATP-synthase inhibition triggers a BDQ-tolerant state with increased dependence on glycolysis, anaplerotic flux and PPP-connected central metabolism(13, 33), produced only modest decreases in lower EMP/TCA intermediates (PEP, PYR and CIT) while largely preserving R5P, S7P, FBP, GAP and glycerone-P. In contrast, TBI-166 alone primarily drove PPP-node remodeling, with R5P and S7P increasing and FBP and glycerone-P decreasing, consistent with attenuated EMP/TCA flux and diversion of G6P/F6P into the PPP. Notably, when BDQ was combined with TBI-166, these patterns shifted from partial remodeling to a more pronounced collapse of BDQ-induced compensatory metabolism: FBP, GAP, PEP, PYR and glycerone-P all fell to significantly lower levels than with either monotherapy, and CIT was further reduced, indicating that, in the presence of ATP-synthase inhibition, TBI-166 prevents MTB from sustaining BDQ-driven glycolytic and anaplerotic input into the TCA/glyoxylate node(13, 31). At the same time, PPP intermediates, particularly R5P and S7P, accumulated more strongly than with TBI-166 or BDQ alone, indicating that carbon is increasingly trapped within PPP-dependent nucleotide and redox metabolism rather than feeding NADH-generating EMP/TCA reactions.

The addition of PZA to the TBI-166+BDQ regimen further accentuated these trends. In the triple-drug group, EMP/TCA intermediates (FBP, GAP, PEP, PYR, glycerone-P and CIT) reached their lowest levels, whereas R5P and S7P showed the highest accumulation across all treatment conditions (Figure 3B). Given that BDQ tolerance involves a coordinated program of enhanced glycolysis, trehalose-fuelled central carbon metabolism and stress-responsive TCA/glyoxylate rerouting to sustain ATP and NADPH under ATP synthase inhibition(31, 34, 35), these quantitative CCM data indicate that TBI-166 metabolically disables this BDQ tolerance program by blocking entry points into the TCA and glyoxylate cycles and depleting the NAD(H) pool. PZA then amplifies this effect, yielding a state in which oxidative phosphorylation and EMP/TCA-linked substrate-level phosphorylation are jointly constrained, while carbon is forced into non-energy-generating PPP and amino-acid/NAD⁺ biosynthetic routes(13, 36, 37). This multilayered disruption of BDQ-induced metabolic remodeling provides a pathway-level explanation for the superior bactericidal activity of the TBI-166+BDQ+PZA regimen.

### 3.4 TBI-166 Suppresses the DosR Dormancy Regulon and Potentiates Triple-Drug Activity against Non-replicating *Mycobacterium tuberculosis*

To further elucidate the mechanism of action of TBI-166 and link its bioenergetic effects to stress adaptation, we first performed transcriptomic profiling of replicating H37Rv exposed to TBI-166(5×MIC). The volcano plot revealed that most significantly downregulated genes belonged to the DosR dormancy regulon, whereas only a small subset of genes was modestly upregulated (Figure 4A). Activated via phosphorylation by the heme-containing sensor kinase DosT (*Rv2027c*), DosR facilitates MTB transition into NR-MTB under low oxygen or elevated levels of H_2_S, NO, and CO (38–42).Consistently, the clustered heatmap showed coordinated repression across multiple DosR regulon blocks, including Rv0079–Rv0081, the hypoxia-responsive cluster Rv1733c–Rv1738 (narX, narK2), Rv1996–Rv1998c and Rv2003c–Rv2007c (fdxA), the pfkB-containing block Rv2028c–Rv2032, the persistence-associated block Rv2623–Rv2631, and the dosR/tgs1-containing block Rv3126c–Rv3134c (Figure 4B). These loci normally support adaptation to low-oxygen and nitrosative stress by enabling nitrate respiration, maintaining NADH/NADPH redox balance and promoting a low-ATP, non-replicating state. Their concerted downregulation indicates that short-term TBI-166 exposure acutely suppresses the DosR dormancy program, suggesting that the ability of MTB to enter or maintain a protected non-replicating phenotype is compromised (43, 44).

**Figure 4.**
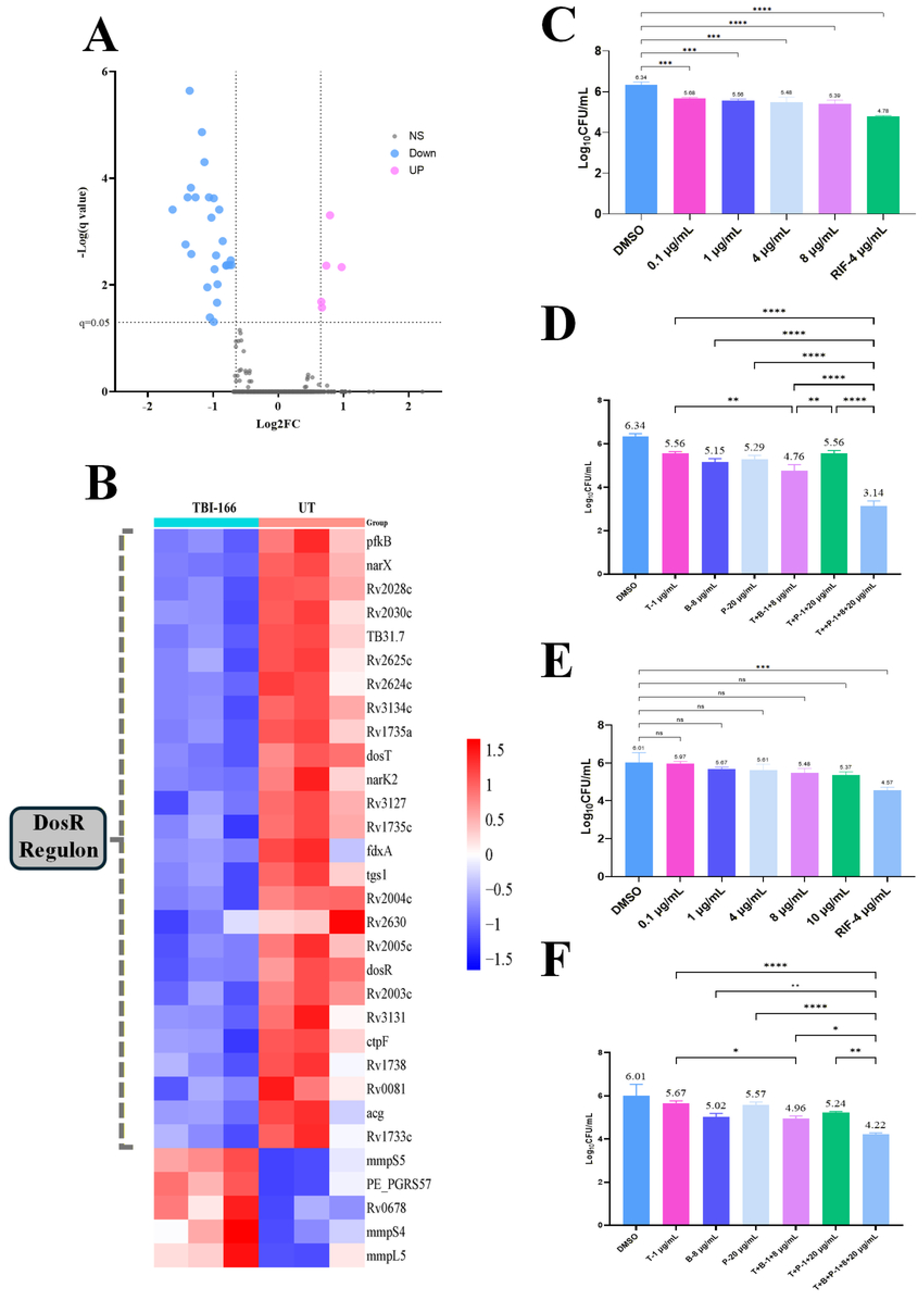
TBI-166 supp.-esses the DosR regulon and enhances TBI-166-based triple efficacy against non-1·eplicating *Mycobacterium tuberculosis*. (A)Volcano plot of differentially expressed genes in replicating MTB treated with TBI-166 (5 X MIC)for 4 hours. The vertical dashed lines indicate the Log_2_FC thresholds of± 0.65, the horizontal dashed line indicates the significance threshold (q = 0.05). Pink dots (UP): genes with significant upregulation. defined as Log_2_FC *:::.* +0.65 and q ≤ 0.05.Blue dots (DOWN): genes with significant downregulation.defined as Log_2_FC:::-0.65 and q:::0.05.Grey dots (NS): genes not statistically significant. (B) Clustered heatmap showing coordinated expression changes of the differentially expressed genes set. The color gradient reflects the fold change (FC) relative to the Untreated(UT). The results are based on three biological replicates (n = 3). (C-D) CFU counts of starvation-induced non-replicating(NR) MTB treated with TBI-166(C) and the combination of TBI-166. BDQ, PZA(D). (E-F) CFU counts of Wayne-induced NR MTB treated with TBI-l66(E) and the combination (F). T: TBI-166: B: BDQ: P: PZA;RIF:Rifampicin. ns: p>0.05: *:p<0.05; **: p<0.01: ***p<0.001; ****: p<0.0001.

Because DosR is a key regulator of non-replicating survival, we next examined whether TBI-166-mediated repression of the DosR regulon enhances TBI-166-based regimen activity against non-replicating bacilli. In the starvation-induced non-replicating model, TBI-166 monotherapy significantly reduced CFU compared with the DMSO control, and combining TBI-166 with either BDQ or PZA further decreased bacterial burden. The TBI-166+BDQ+PZA triple regimen produced the greatest killing, reducing CFU to near the limit of detection (Figure 4C, 4D). In the hypoxia-induced Wayne model, TBI-166 alone exhibited only modest activity, similar to clofazimine; however, addition of BDQ and PZA converted this partial inhibition into strong bactericidal activity, with the triple combination again outperforming all single and dual regimens (Figure 4E, 4F). Across both non-replicating models, the rank order of activity was TBI-166+BDQ+PZA > TBI-166+BDQ > TBI-166+PZA, and inclusion of TBI-166 and PZA consistently accelerated BDQ-mediated CFU decline. Taken together with the ATP depletion, ROS overproduction and central-carbon rewiring observed in replicating bacilli, these findings indicate that TBI-166 simultaneously disrupts energy and redox metabolism and suppresses the DosR-dependent dormancy program, thereby weakening metabolic flexibility and sensitizing non-replicating MTB to the TBI-166+BDQ+PZA combination.

### 3.5 Proteomic Identification of DlaT and LpdC as Candidate Targets of TBI-166, Followed by Multi-Layered Biochemical, Biophysical, and Genetic Validation

To systematically identify the molecular targets through which TBI-166 disrupts CCM and redox homeostasis, we employed a DIA-based quantitative proteomic approach. H37Rv was treated with TBI-166 at 2×, 4×, and 8×MIC for 4 and 24 h, and the global proteomic response was profiled. These proteomic results, combined with the functional significance of DlaT (Rv2215) and LpdC (Rv0462) as parts of multiple enzyme complexes involved in redox homeostasis, EMP and TCA cycle metabolism (31) —identified DlaT and LpdC as the primary candidate targets of TBI-166.The proteomic heatmap (Figure 5A) provides a detailed quantitative view of DlaT and LpdC protein abundance changes across all six treatment conditions. Magenta denotes fold-change > 1 (upregulation) and blue denotes fold-change < 1 (downregulation), as indicated by the color scale. DlaT abundance showed an early modest decrease at 4 h followed by a pronounced and dose-dependent reduction at 24 h across all treatment groups, while LpdC levels increased by approximately twofold at both time points.

**Figure 5.**
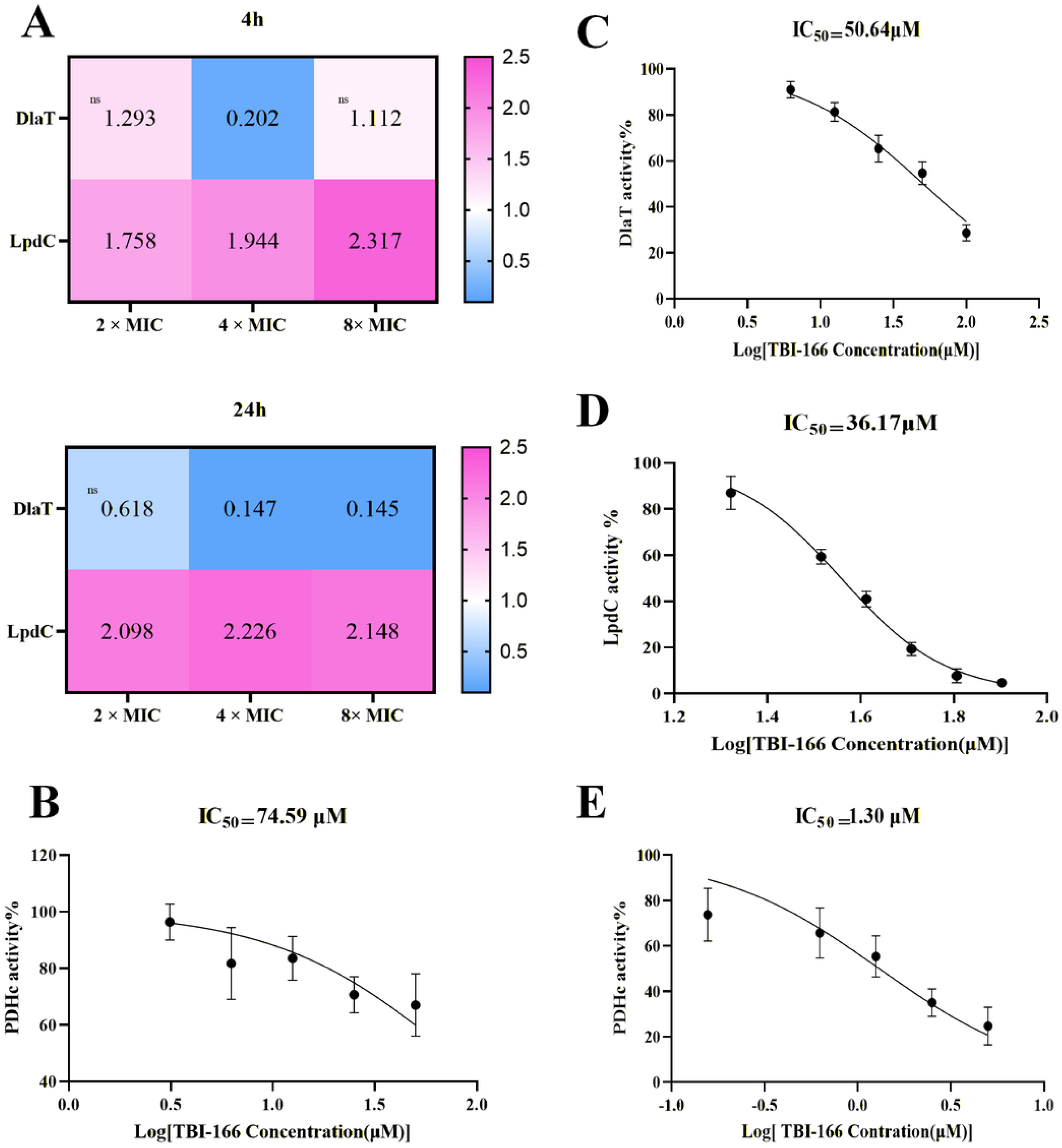
Proteomics implicates DlaT and LpdC as TBI-166 tnrget candidates in *Mycobacterium tuberculosis,* rnlidated by PDHc inhibition assays. (A) Proteomic fold-change heat map showing the relative abundance ofDlaT (Rv2215) and LpdC (Rv0462) upon TBI-166 treatment across the indicated conditions. Magenta and blue indicate fold-change (FC) values relative to the DMSO control. with FC > I shown in magenta and FC< I shown in blue: ns: not significant. (B) Inhibition ofMTB Pyruvate Dehydrogenase Complex (PDHc) crude enzyme activity. (C) Inhibition of purified LpdC enzvme activity. (D) Inhibition of purified DlaT enzyme activity. (E) Inhibition of the reconstituted PDHc (mixture of purified AceE. DlaT, and LpdC). The v-axis represents the remaining enzyme activity (%), normalized lo the DMSO control. Enzyme activity curves are fitted by nonlinear regression.

DlaT and LpdC are key metabolic enzymes in MTB, forming components of multiple enzyme complexes. They participate in the pyruvate dehydrogenase complex (PDHc) with AceE (Rv2241) and the α-ketoglutarate dehydrogenase complex (KDHc) with HOAS (Rv1248c), impacting EMP and the TCA cycle. Furthermore, they contribute to ROS detoxification through peroxidase complex formation with AhpD (Rv2429) and AhpC (Rv2428) within the PNR/P system, maintaining cellular redox balance (45–48). LpdC also participates in branched-chain keto acid dehydrogenase with BkdA (Rv2497c), BkdB (Rv2496c), and BkdC (Rv2495c), influencing protein synthesis(49). Previous studies have established that genetic disruption of DlaT or LpdC severely attenuates MTB virulence in mice and compromises resistance to nitrosative stress, underscoring the essentiality of this enzymatic hub for in vivo persistence(50, 51).

Based on this proteomic lead, we proceeded to direct biochemical validation. We first demonstrated that TBI-166 inhibits MTB PDHc activity in crude lysate supernatants in a concentration-dependent manner, with an IC₅₀ of 74.59 μM (Figure 5B). Using purified recombinant proteins, TBI-166 directly inhibited DlaT and LpdC enzymatic activities with IC₅₀ values of 50.64 μM and 36.17 μM, respectively (Figure 5C, 5D). Notably, when PDHc was fully reconstituted from purified AceE (E1), DlaT (E2), and LpdC (E3), TBI-166 inhibited the complete complex with a markedly lower IC₅₀ of 1.30 μM (Figure 5E)—approximately 30-to 40-fold more potent than against the individual subunits. This sub-micromolar potency against the intact complex suggests that simultaneous engagement of both E2 and E3 within the assembled PDHc architecture produces cooperative functional blockade, consistent with the strong bioenergetic and redox perturbations observed in whole cells.

Biophysical binding studies provided orthogonal confirmation of direct target engagement. Microscale thermophoresis (MST) demonstrated that TBI-166 binds DlaT with a KD of 7.95 ± 2.17 μM and LpdC with a KD of 38.01 ± 15.9 μM, whereas CFZ displayed substantially weaker affinities for both proteins (Table 1) (Figure S3). Differential scanning fluorimetry (DSF) revealed that TBI-166 modestly decreased the melting temperature (Tm) of DlaT from 63.04 ± 0.06 °C to 62.65 ± 0.016 °C, consistent with local destabilization upon ligand binding, while CFZ increased the DlaT Tm. LpdC exhibited very high intrinsic thermostability and was unaffected by TBI-166, but was strongly destabilized by CFZ (Tm reduced to 47.93 ± 0.125 °C) (Table 1) (Figure S2). These distinct biophysical signatures indicate that TBI-166 and CFZ engage DlaT and LpdC in fundamentally different binding modes, with TBI-166 showing superior affinity that matches its stronger PDHc inhibition.

**Table 1.** DSF and MST reveal binding characteristics of TBI-166 with DlaT and LpdC.

| Protein | Small<br>Molecule | MST<br>KD(μM) | DSF<br>Tm($^{\circ}$C) |
| --- | --- | --- | --- |
| DlaT | DMSO | / | 63.04 ± 0.06 |
| | TBI-166 | 7.95 ± 2.17μM | 62.65 ± 0.016 |
| | CFZ | 66.9 ± 26.3μM | 64.38 ± 0.083 |
| LpdC | DMSO | / | ND |
| | TBI-166 | 38.01 ± 15.9μM | ND |
| | CFZ | 50.8 ± 21.6μM | 47.93 ± 0.125 |

/: Not applicable; ND: Not detected. MST provides the dissociation constant (KD, μM), and DSF provides the melting temperature shift (Tm, $^{\circ}$ C). DMSO was used as the vehicle control, CFZ was included as a comparator.

To validate DlaT and LpdC as functionally relevant targets in the cellular context, we performed genetic gain-and loss-of-function experiments. In *M. smegmatis*, CRISPRi-mediated knockdown of either DlaT or LpdC converted the intrinsically resistant wild type (MIC >100 μg/mL) into a hypersusceptible phenotype with MIC values of 0.1953 μg/mL, corresponding to approximately 500-fold sensitization (Figure 7A). In H37Rv, knockdown of DlaT or LpdC similarly reduced the TBI-166 MIC by approximately 4–16-fold compared with wild type (Figure 7B). Conversely, overexpression of either DlaT or LpdC in H37Rv decreased the MIC to roughly one quarter of the wild-type value (Figure 7C). These genetic results are fully consistent with earlier reports that loss of DlaT or LpdC cripples PDHc function, disrupts redox homeostasis, and abrogates virulence in mice(52).

To further distinguish direct target effects from indirect perturbations of associated enzyme complexes, we systematically overexpressed additional PDHc, BCKADH, and PNR/P subunits in H37Rv, including AceE (Rv2241, PDH E1), BkdA/B (Rv2497c/Rv2496c, BCKADH E1/E2), and AhpC/AhpD (Rv2428/Rv2429, PNR/P components). In contrast to the dramatic MIC shifts observed upon DlaT or LpdC modulation, overexpression of these partner proteins only produced modest (±2 log₂-fold) or bidirectional MIC changes (Figure S5), supporting the conclusion that TBI-166 exerts its primary inhibitory effects through direct engagement of the DlaT–LpdC catalytic core rather than through collateral effects on associated complexes.

In summary, the convergent evidence from unbiased proteomic discovery, targeted biochemical inhibition assays, biophysical binding measurements, and systematic genetic validation—including complementary knockdown, overexpression, and partner-protein modulation studies—establishes DlaT and LpdC as the direct functional targets of TBI-166. Through simultaneous inhibition of this central metabolic–redox hub, TBI-166 disrupts PDHc-mediated carbon entry into the TCA cycle, compromises BCKADH-dependent branched-chain amino acid metabolism, and impairs the PNR/P antioxidant defense system, collectively explaining the multifaceted impairment of central carbon metabolism and redox homeostasis observed in earlier sections.

### 3.6 Molecular Docking Reveals the Structural Basis of TBI-166 Binding to DlaT and LpdC, Comparative Analysis with Clofazimine

To elucidate the structural basis of TBI-166’s interaction with its newly validated targets, we performed molecular docking using the recently solved cryo-EM structures of mycobacterial DlaT and LpdC. DlaT was recently shown to assemble into a unique hexameric E2p core that serves as the functional scaffold of mycobacterial PDHc, with each trimer presenting a β-propeller domain that mediates trimer–trimer interactions. The availability of both CoA-bound DlaT trimer (PDB: 9Y7V) and DlaT hexamer (PDB: 9Y6T) structures, together with the MTB LpdC structure (PDB: 7KMY), provided a unique opportunity to explore how TBI-166 engages these targets at atomic resolution(50).

In the CoA-bound DlaT trimer (PDB: 9Y7V), TBI-166 was predicted to occupy the CoA-binding pocket, forming hydrogen-bond interactions with Asp416, Leu421, and Tyr521—residues located within or immediately adjacent to the substrate-binding region (Figure 6A). Given that lipoylated DlaT serves as the acyltransferase core that accepts and transfers pyruvate-derived acetyl groups during PDHc catalysis, occupation of the CoA pocket by TBI-166 would be expected to directly block substrate channeling and acetyl-CoA production, consistent with the potent PDHc inhibition observed in reconstituted enzyme assays (IC₅₀ = 1.30 μM; Figure 5E). Docking against the DlaT hexamer (PDB: 9Y6T) further revealed that TBI-166 can bind at the trimer–trimer interface, where it interacts with Lys495 and Glu507—two residues positioned near the structurally defined interface that maintains hexameric assembly (Figure 6B). Notably, the trimer–trimer interface of DlaT is mediated by a unique antiparallel β-sheet interaction involving Ser508 and Ile509, and disruption of this interface impairs PDHc activity. The predicted binding of TBI-166 at this interface suggests that, in addition to competing with CoA at the catalytic pocket, the drug may also interfere with the oligomeric integrity of the DlaT core.

**Figure 6.**
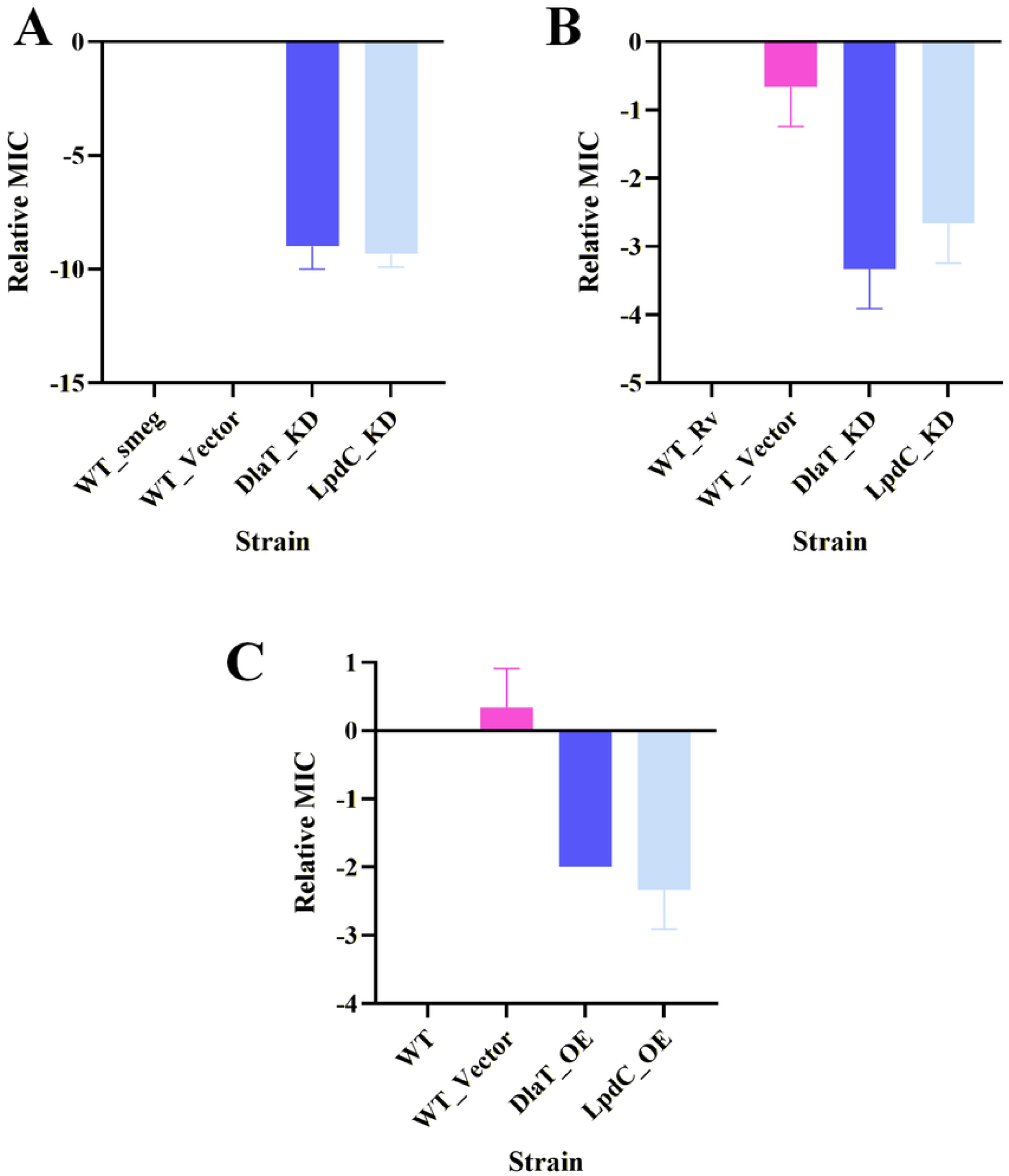
Genetic validation of DlaT and LpdC as targets of TBl-166. (A) Effect of DlaT(Rv2215) and LpdC(Rv0462) knockdown(KD) on susceptibility of *M.smegmatis mc^2^155* to TBI-166. Inducible knockdown strains displayed a marked hypersusceptibility. with the MIC decreasing from> 100 pg/mL in the control strain to 0.09-0.39 pg/mL. (B) Effect of DiaI and LpdC knockdom1 on susceptibility ofH37Rvto TBI-166. Knockdown of either gene decreased the MIC by approximately 4-16 fold compared with wild type. (C) Effect of DiaI and LpdC overexpression on susceptibility ofH37Rv to TBI-166. Overexpression of DlaT or LpdC reduced the MIC to approximately· one quarter of the wild-type value. Bars show. the fold change in MIC relative to the wild-type strain. plotted as logz(MIC_mut/MIC_WT). Data are presented as mean ± SD from at least three independent experiments.

For LpdC (PDB: 7KMY) (53), TBI-166 was placed within the lipoamide-binding channel, forming predicted hydrogen bonds with Arg93, Lys103, and Phe464 (Figure 6C). These residues overlap with structurally characterized functional or ligand-contact regions reported for LpdC. Notably, Phe269 in LpdC has been implicated in a multi-faceted interaction network that contributes to ligand binding and may influence NAD(H) binding or intramolecular electron transfer—a function essential for the re-oxidation of dihydrolipoamide in both PDHc and the PNR/P antioxidant system. The predicted engagement of LpdC’s lipoamide channel by TBI-166 provides a structural rationale for the observed inhibition of LpdC enzymatic activity (IC₅₀ = 36.17 μM; Figure 5D) and is consistent with the MST binding data (KD = 38.01 ± 15.9 μM; Table 1).

As a comparative docking analysis, CFZ was docked into the same DlaT and LpdC structural sites used for TBI-166 docking (Figure S4). In contrast to TBI-166, CFZ formed fewer predicted hydrogen-bond interactions across all three docking models, with only one key hydrogen-bonding residue identified in each site: Thr466 in the CoA-binding pocket of DlaT (PDB: 9Y7V), Glu507 at the DlaT trimer–trimer interface (PDB: 9Y6T), and Arg93 in the LpdC binding pocket (PDB: 7KMY). This reduced hydrogen-bonding network suggests that CFZ adopts a less extensively anchored binding mode within these functional pockets. The decrease in interacting residues may be attributed to differences in molecular geometry, steric fit, and the distribution or accessibility of hydrogen-bond donor/acceptor groups, which may limit the ability of CFZ to simultaneously engage multiple polar residues within the binding cavities. By comparison, TBI-166 formed broader interaction networks with DlaT and LpdC, including multiple hydrogen-bond contacts in the CoA-binding pocket, the trimer–trimer interface, and the LpdC ligand-binding channel. These findings suggest that TBI-166 may achieve improved pocket complementarity and more stable target engagement than CFZ, providing a structural rationale for its potential advantage in interacting with PDHc-associated targets.

Together, these docking results demonstrate that TBI-166 engages both DlaT and LpdC through interactions with conserved functional pockets and interfaces, with a substantially more extensive hydrogen-bonding network than CFZ. For DlaT, the dual engagement of the CoA-binding pocket and the trimer–trimer interface suggests that TBI-166 may interfere with catalytic activity or oligomeric assembly of the PDHc E2 core. For LpdC, occupation of the lipoamide-binding channel provides a mechanism for disrupting the electron transfer cascade within PDHc and the PNR/P system. The comparative analysis with CFZ further highlights the structural basis for TBI-166’s superior target affinity and its multifaceted disruption of central carbon metabolism and redox homeostasis.

## 4. Discussion

One of the key mechanisms underlying MTB drug tolerance is its rapid response to environmental stress (54). Through metabolic downregulation, MTB enters a non-replicating state; while metabolic shifting maintains its homeostasis, enabling successful evasion from drug-mediated killing. This metabolic flexibility poses a significant challenge to the treatment of DR-TB (6), necessitating prolonged therapy and multiple drug combinations to achieve effective cure. However, the high risks of adverse effects, economic burden, and poor patient compliance have driven an urgent need for shorter, all-oral regimens with fewer drugs (55, 56). To address these challenges, drug combinations targeting MTB’s essential metabolic and bioenergetic processes have emerged as promising strategies. Initial efforts focused on targeting multiple branches of the ETC, such as cytochrome bd oxidase, cytochrome bcc-aa3 oxidase, and ATP synthase, which is significantly anti-tuberculosis treatment efficacy(22).

For instance, combinations like CK-2-63+Q203+BDQ demonstrated potent in vitro bactericidal activity(57–59), while the synthetic lethal interaction between Q203 and cytochrome bd inhibitors highlighted the vulnerability arising from respiratory redundancy. More sophisticated approaches have emerged that simultaneously target both ETC and essential metabolic processes. For instance, the synergistic interaction between TB47 (a cytochrome bc1 inhibitor) and pretomanid (PMD) effectively disrupts both ETC function and the PPP(60). Similarly, the BDQ+Q203+CFZ combination exemplifies how targeting respiratory flexibility while generating oxidative stress can transform MTB’s metabolic adaptability into a vulnerability (33). These findings underscore the therapeutic potential of strategies that disable MTB’s adaptive metabolic mechanisms through coordinated multi-target inhibition.

Building on this framework, our work establishes TBI-166 as a prototype “metabolic flexibility breaker” that acts by dual inhibition of DlaT (Rv2215) and LpdC (Rv0462)—the E2 and E3 components of the pyruvate dehydrogenase complex (PDHc)—and extends its impact through a TBI-166+BDQ+PZA regimen. Multi-layered evidence from proteomics, enzymology, biophysics, docking, and genetics consistently converges on the DlaT–LpdC axis as the primary functional target of TBI-166. DlaT and LpdC form the catalytic core of three multienzyme complexes—PDHc, branched-chain α-keto acid dehydrogenase (BCKADH), and the peroxynitrite reductase/peroxidase (PNR/P) system—jointly controlling carbon entry into the TCA cycle, branched-chain amino acid catabolism, and detoxification of reactive nitrogen species(31, 53, 61). By inhibiting DlaT and LpdC, TBI-166 simultaneously dampens PDHc- and BCKADH-dependent flux, leading to pyruvate accumulation and slowing of both the EMP and the TCA cycle, while compromising PNR/P-mediated redox defense. This coordinated blockade collapses the normal buffering capacity of the NAD(H) pool and ATP-generating networks that MTB would ordinarily mobilize in response to BDQ- or PZA-induced stress.

Consistent with this model, global metabolite profiling and ¹³C isotopic tracing (Figures 2 and 3) reveal that in the presence of TBI-166, MTB attempts to compensate by rerouting carbon through alternative pyruvate-linked pathways and TCA cycle branches, but these rerouting efforts remain incomplete and cannot restore metabolic homeostasis. At the transcriptional level, TBI-166 downregulates the DosR dormancy regulon and its downstream effectors (Figure 4), thereby directly weakening a major regulatory program that enables the bacterium to shift into metabolically quiescent, drug-tolerant states. On the protein and enzyme-activity level, proteomic changes and biochemical assays (Figure 5) demonstrate that TBI-166 suppresses PDHc activity and destabilizes the DlaT–LpdC hub, while genetic perturbation of DlaT and LpdC modulates TBI-166 susceptibility (Figure 6). Together, these data support a model in which TBI-166 acts at the intersection of central carbon metabolism and redox homeostasis to restrict the “degrees of freedom” available for metabolic rewiring, thus functionally constraining MTB’s metabolic flexibility.

At the structural level, TBI-166’s ability to break metabolic flexibility is underpinned by its favorable engagement of DlaT and LpdC. DlaT in MTB assembles into a unique concentration-dependent hexameric E2p core that differs fundamentally from the canonical octahedral/icosahedral E2 architectures found in other bacteria(50). Molecular docking suggests that TBI-166 occupies the CoA-binding pocket or the trimer–trimer interface of this hexameric E2p core, while also binding within the lipoamide channel of LpdC (Figure 7). These dual-site interactions provide a structural explanation for the potent inhibition of the reconstituted PDHc complex, and for the tighter in vitro binding affinities of TBI-166 to DlaT/LpdC compared with clofazimine. Importantly, by targeting a structurally distinctive E2p core and an enzymatic hub absent from human mitochondria in this configuration, TBI-166 exploits a mycobacteria-specific vulnerability at the heart of CCM and redox control.

**Figure 7.**
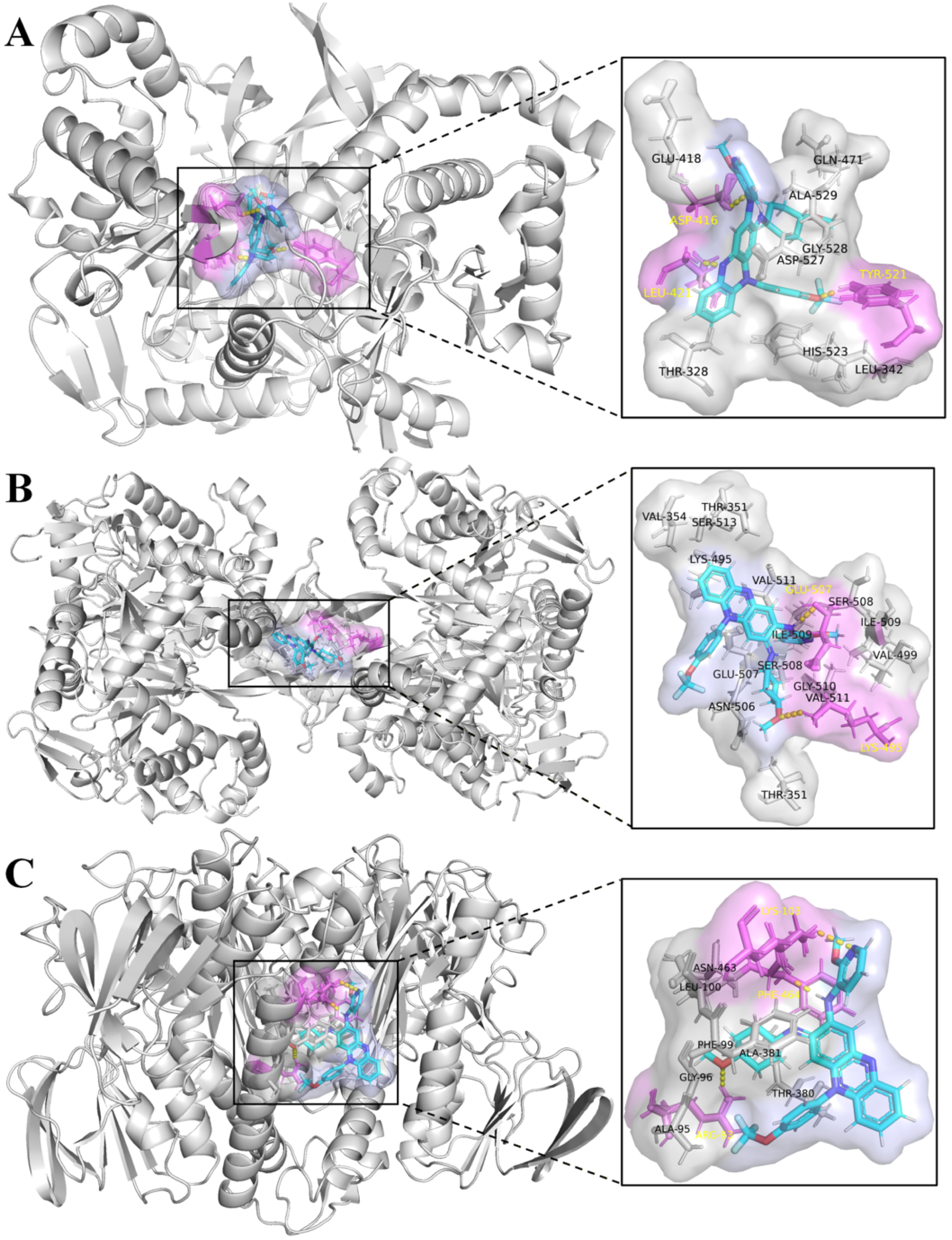
Molecular docking predicts Pyrifazimine binding to DlaT and LpdC in Mycobncterium tuberculosis. (A) Molecular docking of Pyrifazimine into the CoA-binding pocket of DlaT (R\’2215; PDB ID: 9Y7V). showing predicted hydrogen-bond interactions \\ith ASP-416. LEU-421. and TYR-521. (B) Molecular docking of Pyrifazimine at the trimer-trimer interface of DlaT (PDB ID: 9Y6T). showing predicted hydrogen-bond interactions with LYS-495 and GLU-507. (C) Molecular docking of Pyrifazimine with LpdC (PDB ID: 7KMY). showing predicted hydrogen-bond interactions with ARG-93. LYS-103. and PHE-464. Close-up view show the ligand-binding pockets and interacting residues. Key amino acid residues forming hydrogen-bond interactions with Pyrifazimine are highlighted in purple. and their residue names are labeled in yellow.

Within the TBI-166+BDQ+PZA regimen, this DlaT/LpdC-centered mechanism reshapes how BDQ and PZA act on MTB. BDQ inhibits F₁F₀-ATP synthase and forces the bacterium to rely more heavily on substrate-level phosphorylation and anaplerotic reactions; PZA exerts context-dependent activity that is enhanced when redox and energy homeostasis are already destabilized. By pre-emptively disabling the DlaT–LpdC hub, TBI-166 limits the rerouting options available to sustain ATP production and NAD(H) balance in response to BDQ, and simultaneously weakens detoxification systems and pH/redox buffering that influence PZA susceptibility. As a result, the three drugs cooperate to collapse both respiratory and metabolic plasticity across aerobic replicating and non-replicating states, leading to strong early bactericidal and sterilizing effects in murine models. Conceptually, TBI-166 transforms MTB’s normally protective metabolic adaptability into a liability that can be exploited by BDQ and PZA.

From a translational perspective, TBI-166 further stands out as a clinically promising riminophenazine analog. Compared with CFZ, TBI-166 maintains comparable antimycobacterial activity but shows reduced skin pigmentation and favorable interaction with BDQ by lowering toxic BDQ metabolite formation(1, 17). Coupled with its unique DlaT/LpdC-targeting profile and ability to constrain metabolic flexibility, these pharmacodynamic and safety attributes provide a compelling rationale for prioritizing TBI-166-containing, metabolism-centered regimens in DR-TB development pipelines. More broadly, this work illustrates that directly targeting metabolic hubs that coordinate CCM, redox homeostasis, and dormancy regulation can convert drug-tolerant metabolic states into drug-susceptible ones, suggesting a generalizable design principle for future regimens.

Several limitations warrant consideration. First, although our mechanistic conclusions are supported by multi-omic, biochemical, structural, and genetic data, they are primarily derived from in vitro systems and murine models, which do not fully capture the microenvironmental heterogeneity of human TB lesions. Second, while our data link DlaT/LpdC inhibition to NAD(H) imbalance and DosR repression, the precise causal circuitry connecting these events remains to be fully resolved; additional targets or signaling pathways may contribute to the observed phenotypes. Third, we have not yet systematically dissected how simultaneous modulation of DlaT and LpdC—for example, double knockdown or co-overexpression—affects TBI-166 activity and the global metabolic state, which will be important to definitively establish the quantitative contribution of each target to metabolic flexibility. Finally, the predicted binding modes of TBI-166 and CFZ on DlaT/LpdC are based on computational docking and require experimental confirmation by structural biology and high-resolution binding studies. Addressing these gaps will refine our understanding of how DlaT/LpdC-centered interventions can be optimized to maximally disrupt MTB metabolic adaptability.

## 5. Conclusion

In summary, this study identifies TBI-166 as a central multitarget agent that breaks M. tuberculosis metabolic flexibility by dual inhibition of DlaT and LpdC at the core of PDHc- and BCKADH-linked central carbon metabolism and the PNR/P redox defense system. By destabilizing this metabolic–redox hub, TBI-166 disrupts NAD(H) homeostasis, downregulates the DosR dormancy regulon (Figure 4), and constrains the bacterium’s capacity to rewire carbon flux in response to stress. When combined with BDQ and PZA, which target oxidative phosphorylation and stress-dependent vulnerabilities, TBI-166 converts MTB’s normally protective bioenergetic and metabolic plasticity into a point of fragility. The TBI-166+BDQ+PZA regimen thus achieves potent bactericidal and sterilizing activity against both replicating and non-replicating MTB by simultaneously collapsing energy metabolism, redox homeostasis, and adaptive metabolic remodeling. These findings support a strategy for DR-TB therapy in which direct targeting of metabolic flexibility—exemplified by the DlaT–LpdC axis—serves as the mechanistic backbone of short, fully oral combination regimens.

## Acknowledgement

The authors declare that the research was conducted in the absence of any commercial or financial relationships that could be construed as a potential conflict of interest.

This work was supported by the National Natural Science Foundation of China (82173862) and Beijing Hospitals Authority’s Ascent Plan Support (DFL20221402).

During the preparation of this work the authors used Chatgpt and DeepSeek in order to language polishing. After using this tool, the author(s) reviewed and edited the content as needed and takes full responsibility for the content of the publication.

